# Withaferin A promotes white adipose browning and prevents obesity through sympathetic nerve-activated Prdm16-FATP1 axis

**DOI:** 10.1101/2021.02.25.432705

**Authors:** Bingbing Guo, Jiarui Liu, Bingwei Wang, Chenyu Zhang, Zhijie Su, Miao Zhao, Ruimao Zheng

## Abstract

The increasing prevalence of obesity has resulted in demands for the development of new effective strategies for obesity treatment. The Withaferin A (WA) shows a great potential for prevention of obesity by sensitizing leptin signaling in the hypothalamus. However, the mechanism underlying the weight- and adiposity-reducing effects of WA remains to be elucidated. Here, we report that WA treatment induced white adipose tissue (WAT) browning, elevated energy expenditure (EE), decreased respiratory exchange ratio (RER), and prevented high-fat diet (HFD)-induced obesity. The sympathetic chemical denervation dampened the WAT browning and also impeded the reduction of adiposity in WA-treated mice. WA markedly up-regulated the levels of Prdm16 and FATP1 (Slc27a1) in the inguinal WAT (iWAT), and this was blocked by sympathetic denervation. Prdm16 or FATP1 knockdown in iWAT abrogated the WAT browning-inducing effects of WA, and restored the weight gain and the adiposity in WA-treated mice. Together, these findings suggest that WA induces WAT browning through the sympathetic nerve-adipose axis; and the adipocytic Prdm16-FATP1 pathway mediates the promotive effects of WA on white adipose browning.

## Introduction

Obesity is defined as a medical condition of abnormal or excessive fat accumulation in adipose tissue, to the extent that the energy homeostasis is disrupted, which has become a major risk factor for the onset and progression of type 2 diabetes, nonalcoholic fatty liver disease and cardiovascular disease (González-Muniesa et al., 2017; Williams et al., 2015). Over the last 30 years, the prevalence of obesity in many countries has doubled, or even quadrupled, fueled by excessive food intake, lackness of physical activity, and adaptive genetic variation of human that facilitates energy storage (Qasim et al., 2018; Ramachandrappa and Farooqi, 2011). To date, obesity has been one of the leading preventable causes of death worldwide (Bauer et al., 2014). However, the effective pharmacotherapy for obesity is still lacking.

Withaferin A (WA) is a steroidal lactone derived from Withania somnifera, a traditional medicinal herb prescribed for a variety of ailments owing to its anti-inflammatory, anti-tumor, anti-diabetic, and neuroprotective properties (Dutta et al., 2019). WA is highly lipid-soluble and can cross the blood-brain barrier (BBB) (Zahiruddin et al., 2020). Recently, it is reported that WA enhances leptin sensitivity and prevents obesity by targeting hypothalamic neurons. Leptin is a key adipokine that plays crucial roles in the maintenance of energy balance and metabolic homeostasis (Pan and Myers, 2018). Leptin activates its signal pathway in hypothalamus, leading to increased fat utilization, elevated energy expenditure (EE), promoted white adipose tissue (WAT) browning, and counteracts obesity development (Caron et al., 2018; Schwartz et al., 2000). The elevated plasma level of leptin coexists with the excessive adiposity in obese human and rodents, which is interpreted as evidence of leptin resistance (Myers et al., 2008). Leptin resistance impedes the clinical application of leptin for obesity treatment (Cardinali and Hardeland, 2017; Myers et al., 2012). Thus, sensitizing the leptin signaling to prevent obesity has attracted widespread attention, proposing a novel strategy for obesity treatment. WA potentiates leptin signaling in ARC (arcuate nucleus), DMH (dorsomedial hypothalamic nucleus) and VMH (ventromedial hypothalamic nucleus) to reduce fat mass, and protects against HFD-induced obesity in mice (Lee et al., 2016). However, the mechanism underlying the adiposity-reducing effects of WA remains incompletely understood.

Beige adipocytes are brown-like adipocytes dispersed throughout the white fat depots, which have the capacity for uncoupled respiration and heat production (Wang and Seale, 2016). Stimulation of the beige fat biogenesis in WAT (also known as ’browning’) confers beneficial effects on metabolism, leading to elevated EE, increased thermogenesis and reduced adiposity (Bartelt and Heeren, 2014). In humans and rodents, the WAT depots show potential for the browning, although this potential is weakened in older and obese subjects (Herz and Kiefer, 2019; Sheng et al., 2020). Emerging evidence shows that pharmacological agents, environmental stimuli, or physical exercise can induce WAT browning and reduce body weight (Kim et al., 2017a; Shin et al., 2017; Song et al., 2018). The hypothalamus, as a center for controlling energy balance and metabolic homeostasis, is tightly involved in the regulation of WAT browning through sympathetic nervous system (SNS) (López et al.,

2016; Sladek et al., 2015). Activation of leptin signal in the hypothalamus leads to an increased sympathetic nervous activity (SNA) and an enhanced WAT browning (Dodd et al., 2015). WA potentiates hypothalamic leptin signaling and reduces adiposity (Lee et al., 2016) However, it is unknown whether the sympathetic nerve-mediated WAT browning contributes to the anti-obesity effect of WA.

To explore the neural and molecular mechanisms by which WA induces WAT browning and reduces adiposity, the chemical denervation, genome-wide deep-sequencing analysis, lipidomics analysis, and in vivo RNA interference technology were employed in this study. Our findings identified that WA could be a promising agent for the pharmacotherapy of obesity, and provided a therapeutically useful way of administering WA for treatment of obesity and associated comorbidities.

## Results

### WA promotes WAT browning and prevents diet-induced obesity

To explore the mechanism underlying the anti-obesity effects of WA, we first examined whether WA reduces body weight or adiposity by promoting WAT browning. We found that WA treatment at dosage levels of 2, 20 or 200 μg/kg for 7, 14 or 21 days reduced body weight and fat pad weights, without altering food intake in the high-fat diet (HFD)-fed mice (**Figure S1A-S1L**). WA at dosage levels of 2, 20 or 200 μg/kg led to an increased expression of the browning-associated genes in the iWAT (**Figure S1M-S1O**). Thus, WA treatment at dosage level of 2 μg/kg was selected for further study. We performed WA treatment (2 μg/kg intraperitoneally (ip) once-daily for 2 months) to examine the long-term effects of WA (**Figure 1A**). We found that WA treatment led to a significant and sustained reduction in body weight, equivalent to a total weight loss of 16.1% at week 9 (**Figure 1B**). WA reduced the weights of inguinal, epididymal and perirenal fat pads, but did not affect food intake and the weight of brown adipose tissue (BAT) (**Figure 1C, D**). WA increased expression levels of the browning-associated genes in iWAT (**Figure 1E**). Notably, WA reduced body weight and increased browning-associated gene expressions at as early as week 2 (**Figure 1B, E**). Therefore, the intraperitoneal injection of WA at 2 μg/kg for 7 days was selected for further study. The thermoneutrality (30°C for mice) is critical for examining the pro-browning effect of chemical compounds; thus, we performed the experiments at either room temperature or thermoneutrality (**Figure 1F**). WA reduced body weight and fat pad weights at either room temperature or thermoneutrality, with no change in food intake (**Figure 1G-J**). The canonical adipose browning phenotypes such as “brown-like” adipocytes with smaller and multilocular lipid droplets in iWAT were observed in WA-treated mice (**Figure 1K**). WA increased the expression levels of the genes involved in the WAT browning, the sympathetic nervous activity, and the adaptive thermogenesis in iWAT, BAT and eWAT (**Figure 1L**). The expression levels of UCP1 and PGC1α were significantly elevated in iWAT (**Figure 1M; Figure S2A**). The thermogenesis of iWAT and BAT was increased during the light phase (when mice were inactive) in WA-treated mice (**Figure 1N**). WA led to enhanced whole-body EE and decreased respiratory exchange ratio (RER) (**Figure 1O-P; Figure S3A**). WA did not affect the daily locomotor activity (**Figure S3B**). Together, these findings show that WA induces WAT browning, increases sympathetic activity, and reduces body weight, suggesting that WA may prevent HFD-induced obesity by promoting WAT browning; and the sympathetic activity may contribute to mediating the WAT browning-inducing effects of WA.

**Figure 1.**
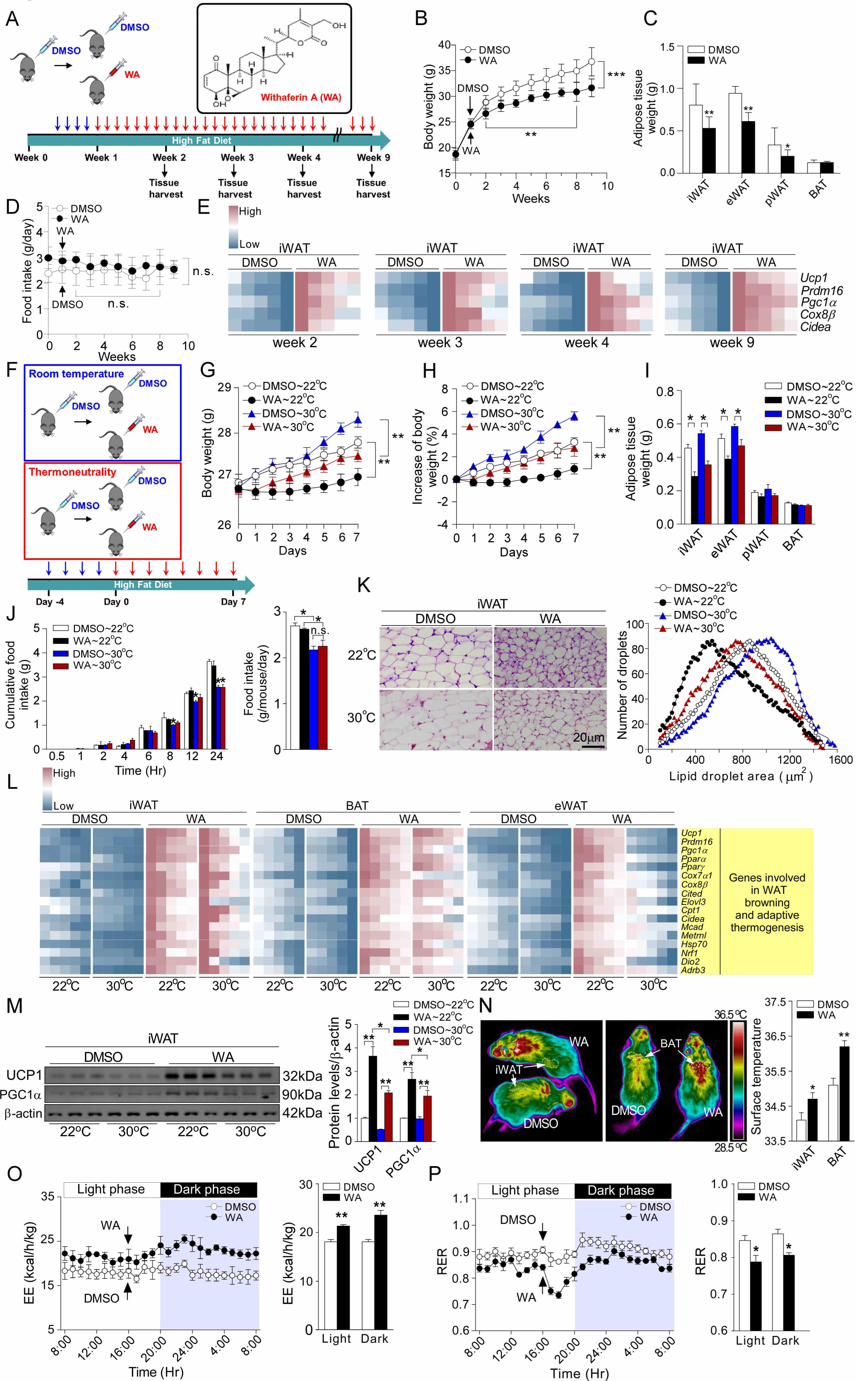
WA promotes WAT browning and prevents diet-induced obesity. **(A-E)** 8 week-old mice were housed at 22°C, fed a HFD and treated with WA or DMSO for 2 months. **(A)** Schematic illustration of experiments. Mice received DMSO for 4 days as acclimation, then were treated with WA or DMSO daily for 2 months. Tissue harvest was performed at week 2, 3, 4 and 9. **(B)** Body weight (WA, n = 6; DMSO, n = 5). **(C)** Adipose tissues weight (WA, n = 6; DMSO, n = 5). **(D)** Daily food intake (WA, n = 6; DMSO, n = 5). **(E)** Heatmap visualization of relative mRNA expression of browning markers in iWAT (n = 5). **(F-P)** 8 week-old mice were housed at either 22°C or 30°C, fed a HFD and treated with WA or DMSO for 7 days (WA∼22°C, n = 7; DMSO∼22°C, n = 6; WA∼30°C, n = 8; DMSO∼30°C, n = 7). **(F)** Schematic illustration of experiments. Mice received DMSO for 4 days as acclimation, then were treated with WA or DMSO daily for 7 days. Metabolic chamber and tissue harvest were performed at day 7. **(G)** Body weight. **(H)** Body weight gain. **(I)** Adipose tissues weight. **(J)** Cumulative food intake. **(K)** Left panel, H&E staining. Scale bar indicates 20 μm. Right panel, cell size profiling of adipocytes from iWAT. **(L)** Heatmap visualization of Relative mRNA expression of browning-associated genes in iWAT, BAT and eWAT (n = 6). **(M)** Western blot with quantification of Ucp-1 and PGC1α in iWAT. **(N)** Left panel, representative thermographic image of WA and DMSO at 22°C; Right panel, calculated averages of iWAT and BAT surface temperature (n = 5). (**O-P**) Indirect calorimetry was performed to quantify EE (**Q**) and RER (**R**) of WA or DMSO-treated mice at 22°C during complete 24 hr light-dark cycles, the arrow indicates time of WA or DMSO injection (n = 5). Values were expressed as mean ± SEM. Statistical analysis: Student’s t test for **C**, **E**, **N**, **O-P**; two-way ANOVA for **B**, **D**; and one-way ANOVA with Bonferroni test for the rest. *p < 0.05, **p < 0.01, and ***p < 0.001; n.s., not significant.

### Sympathetic nerve-adiposity connection contributes to WA-induced WAT browning

To determine the role of sympathetic nerves in mediating the WAT browning-inducing effects of WA, we assessed the leptin signal activity in the hypothalamus and examined the sympathetic nervous activity in iWAT. We found that WA treatment upregulated the gene expressions associated with activation of classic leptin signal pathway (LepR, JAK2, Stat3/5, PI3K, AKT, mTOR, S6K1, PDE3b, AMPK, Mchr1 and Pomc); and also increased expression levels of the positive regulators of leptin signaling (Rock1, Sitr1, MFN2 and BDNF) (**Figure 2A**). The tyrosine hydroxylase (TH), a key synthase of norepinephrine, and an indicator of the activity of sympathetic nerves, was significantly increased in iWAT treated with WA (**Figure 2C-D; Figure S4A**). Of note, the unilateral sympathetic denervation of iWAT by 6-OHDA restored the size of adipocytes in iWAT, as compared with contralateral sham-operation (**Figure 2E-F**). Moreover, the WA-increased expression levels of the browning-associated genes were also normalized by unilateral sympathetic denervation (**Figure 2G-H; Figure S4B**). Taken together, these findings indicate that WA enhances leptin sensitivity in the hypothalamus, and the sympathetic innervation is required for the WA-induced WAT browning.

**Figure 2.**
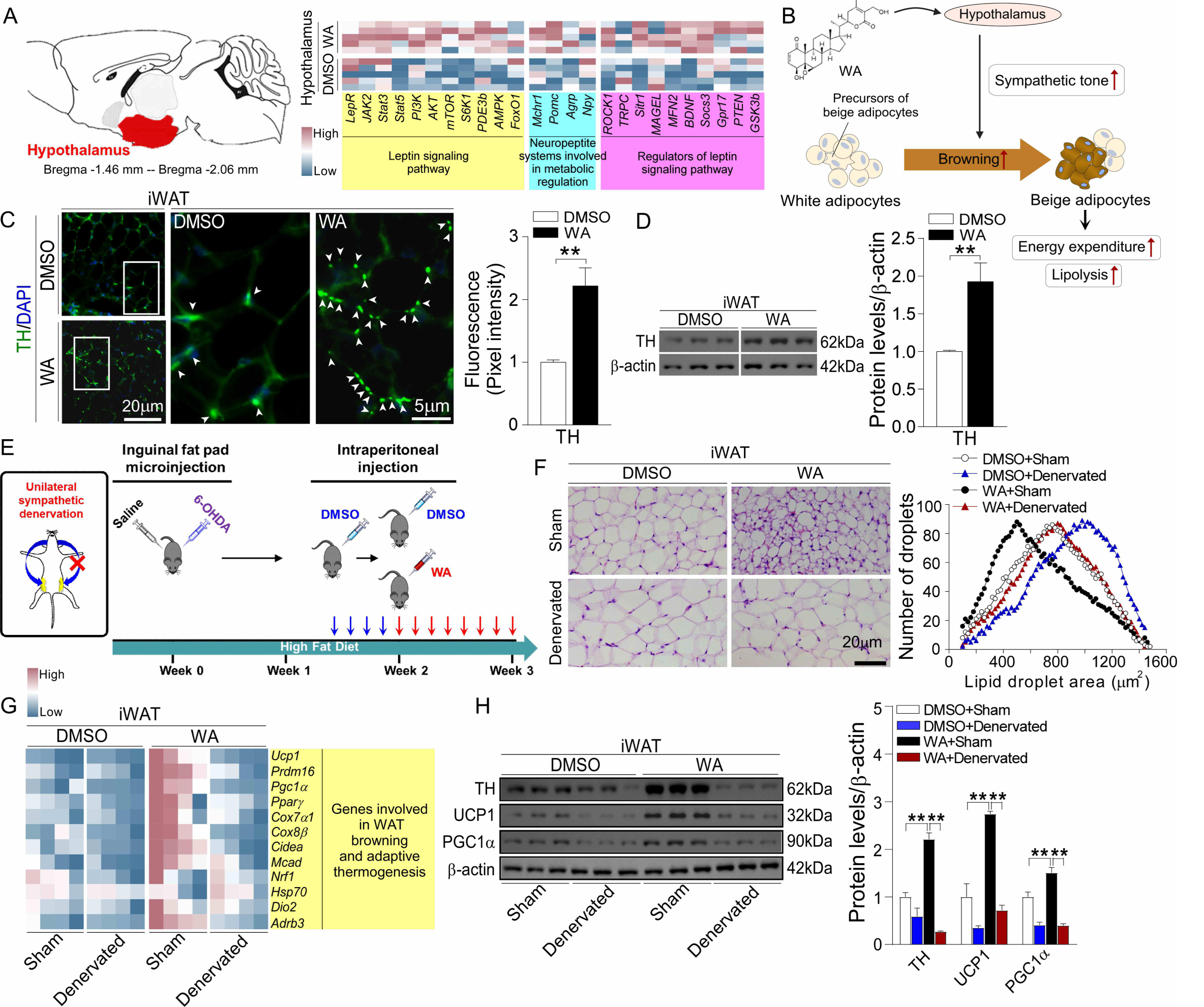
Sympathetic nerve-adiposity connection contributes to WA-induced WAT browning. **(A)** Heatmap visualization of relative mRNA expression of the genes associated with leptin signaling in hypothalamus from mice treated with WA or DMSO (n = 5). **(B)** Schematic illustration showing the regulation of WAT browning by hypothalamus-SNS pathway. **(C)** Immunofluorescence staining of TH in iWAT. Scale bar indicates 20 μm or 5 μm. **(D)** Western blot with quantification of TH in iWAT. The experiments in **A-D** were repeated three times with similar outcomes. **(E-G)** 8 week-old mice were unilaterally denervated (6-OHDA) and then treated with WA or DMSO daily for 7 days (WA, n = 8; DMSO, n = 6). **(E)** Schematic illustration of experiments. The iWAT in 8 week-old mice was unilaterally denervated. 2 weeks later, mice were treated with WA or DMSO daily for 7 days. Tissue harvest was performed at week 3. **(F)** Left panel, H&E staining. Scale bar indicates 20 μm. Right panel, cell size profiling of adipocytes from iWAT. **(G)** Heatmap visualization of relative mRNA expression of WAT browning-associated genes in iWAT (n = 4). **(H)** Western blot with quantification of TH, Ucp-1 and PGC1α in iWAT. Values were expressed as mean ± SEM. Statistical analysis: Student’s t test for **A**, **C**, **D**; and one-way ANOVA with Bonferroni test for **G**, **H**. *p < 0.05, **p < 0.01, and ***p < 0.001.

### WA promotes WAT browning and elevates EE by sympathetic–adipose pathway

To examine the extent to which the sympathetic nerve-mediated WAT browning may contribute to the elevation of EE and the reduction in body weight in WA-treated mice, we performed bilateral denervation of iWAT (**Figure 3A**) (Dodd et al., 2015). We found that the reduction in body weight, the elevation of EE, as well as the decrease in RER were diminished by bilateral denervation of iWATs in WA-treated mice (**Figure 3B-E; Figure S5A**). The motor activity was not affected by bilateral denervation (**Figure S5B**). Notably, WA elevated EE by 10.1% in the bilaterally sympathetic denervated mice, compared with 25.3% in sham-operated mice (**Figure 3D**), which highlights that the activity of sympathetic nerves in white adipose tissue is essential for mediating the obesity-preventive effects of WA. The lipidomic analysis showed that WA treatment increased the levels of oleic and arachidonic acid, and decreased the level of a saturated fatty acid (tricosanoic acid) in iWAT, which were dampened by iWAT bilateral sympathetic denervation (**Figure 3F**), suggesting that WA may modify the fatty acid composition in iWAT via SNS, which may also be implicated in the promotion of the biological process of WAT browning. Taken together, these observations suggest that the sympathetic denervation abrogates the WAT browning by blocking the sympathetic signal from the hypothalamus to the white adipose tissue, and demonstrate that WA-induced WAT browning is mediated by the hypothalamus-SNS-WAT axis.

**Figure 3.**
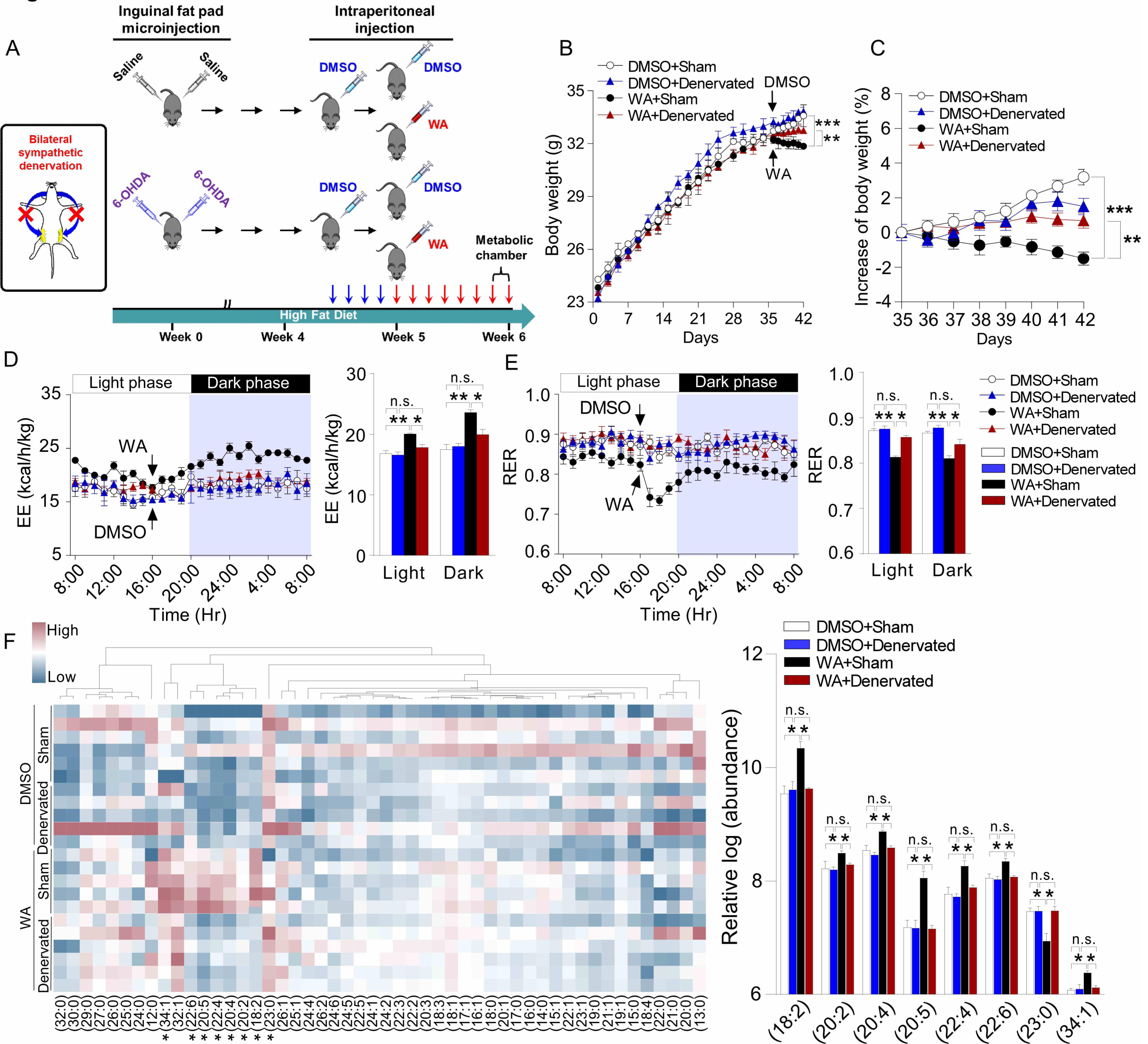
WA promotes WAT browning and elevates EE by sympathetic–adipose pathway. 8 week-old mice were bilaterally denervated (6-OHDA) and then treated with WA or DMSO daily for 7 days (DMSO+Sham, n = 6; DMSO+Denevated, n = 8; WA+Sham, n = 6; WA+Denevated, n = 9). **(A)** Schematic illustration of experiments. The iWAT in 8 week-old mice was bilaterally denervated (6-OHDA). 5 weeks later, mice were treated with WA or DMSO daily for 7 days. Metabolic chamber and tissue harvest were performed at week 6. **(B)** Body weight. **(C)** Body weight gain. **(D-E)** Indirect calorimetry was performed to quantify EE (**D**) and RER (**E**) of denervated or sham-operated mice treated with WA or DMSO during complete 24 hr light-dark cycles, the arrow indicates the time of WA or DMSO injection (n = 5). **(F)** Left panel, lipidomic heatmap showing the average changes in free fatty acids of iWAT; Right panel, relative changes in the significantly altered species of fatty acids in iWAT (DMSO+Sham, n = 6; DMSO+Denevated, n = 5; WA+Sham, n = 5; WA+Denevated, n = 6). Values were expressed as mean ± SEM. Significance was determined by one-way ANOVA with Bonferroni test. *p < 0.05, **p < 0.01, and ***p < 0.001.

### Prdm16 and FATP1 are identified as molecular targets of WA

To dissect the molecular mechanism underlying the WA-induced WAT browning, we performed the genome-wide transcriptomic sequencing analysis for the iWAT in denervated and sham-operated mice. We visualized the gene expression profiles as a heatmap, and observed that WA remarkably increased expression levels of the genes associated with WAT browning (Ucp1, Prdm16, Ppargc1a), mitochondrial biogenesis (Mtfp1, Slc25a25), lipid metabolism (FATP1, Fabp4, Pdk4), glucose metabolism (Sik2, Irs1, Pfkfb1/3), and nervous system function (Negr1, Nr4a3, Trim67, Unc13a, Dok6); while reduced expression levels of the genes associated with immunity (Ccr5, Oas1g, Oas1a, Orm2), neural inhibition (Slc6a2, Mt3) and apoptosis (Cadm1, Gzma) (**Figure 4A-B**). These pleiotropic effects of WA were abrogated by the iWAT sympathetic denervation (**Figure 4A-B**). GO (gene ontogeny) and KEGG (Kyoto Encyclopedia of Genes and Genomes) analyses showed that WA activated the pathways associated with insulin signaling, adaptive thermogenesis and kinase activity regulation; while attenuated the pathway relative to inflammatory response, which were dampened by sympathetic denervation (**Figure 4C-E**). Volcano plot and Venn diagram analysis showed that WA markedly increased the expression levels of FATP1, Ucp1, Prdm16 and Pparγ in iWAT, which were also abolished by sympathetic denervation (**Figure 4F-H**). GeneMANIA prediction showed a potential interaction and crosstalk between Prdm16 and FATP1 in iWAT treated with WA (**Figure 4I**). By analyzing a human subcutaneous adipose tissue dataset, we found that the adipose browning-promoting condition, such as physical exercise, also robustly increased the expression levels of Prdm16 and FATP1, demonstrating that Prdm16 and FATP1 may be important factors in the regulation of WAT browning in human (**Figure 4J**). Taken together, these findings provide a comprehensive insight into the transcriptomic modifications induced by WA, and demonstrate that Prdm16 and FATP1 may be the critical molecular targets in the biological process of WA-induced WAT browning, which contributes to understanding of the potential mechanism by which WA induces WAT browning and prevents obesity.

**Figure 4.**
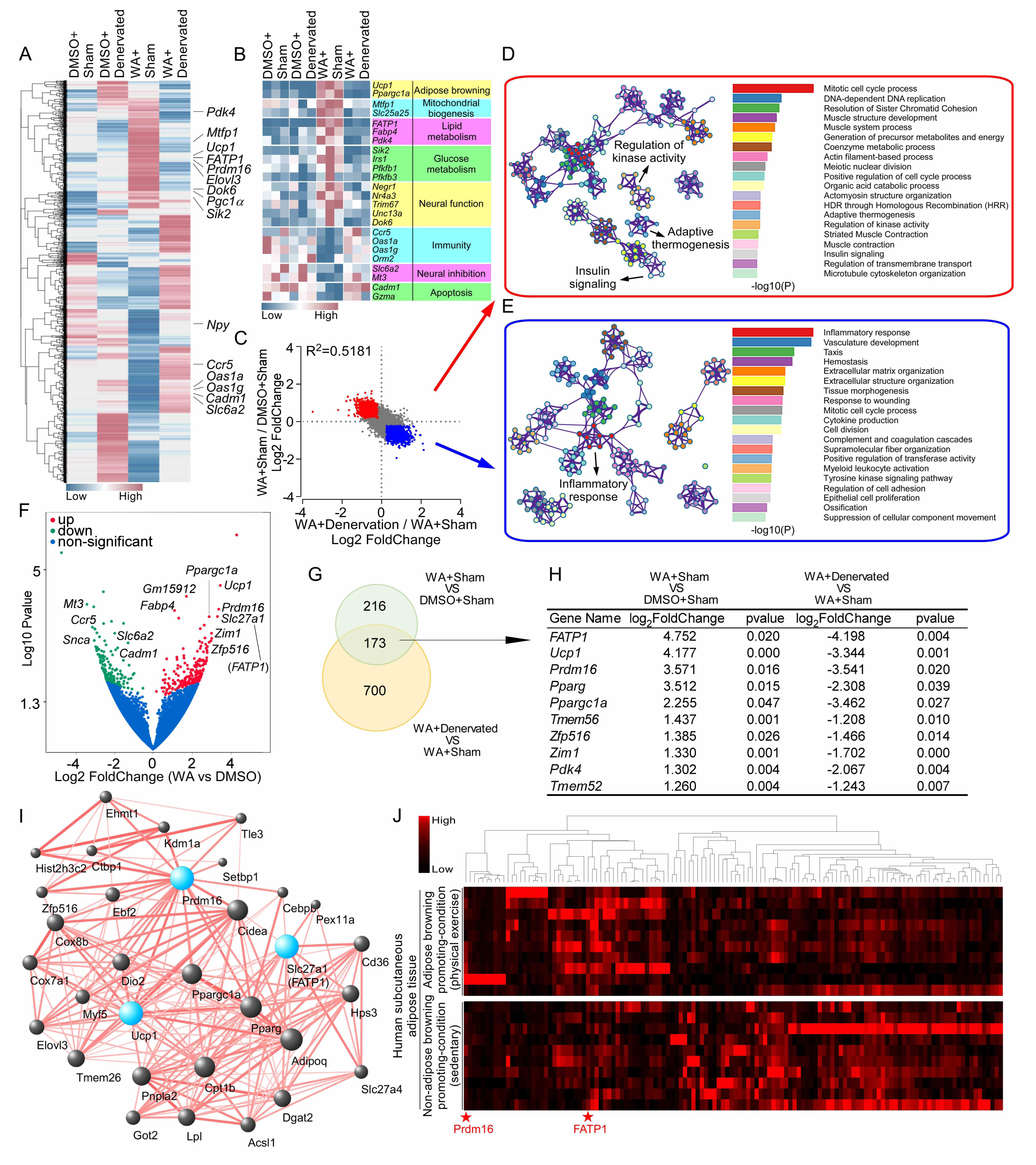
Prdm16 and FATP1 are identified as molecular targets of WA. **(A)** Hierarchical clustered heatmap of gene expression profiles of iWAT of sympathetic-denervated or sham-operated mice in WA or DMSO treatment (n = 3). **(B)** Heatmap of detail relative change in DEGs of iWAT. **(C)**. Scatter plot highlights the genes upregulated by WA and downregulated by sympathetic denervation (Red); and the genes downregulated by WA and upregulated by sympathetic denervation (Blue). **(D, E)** Gene Ontology enrichment based on DEGs that have a P value smaller than 0.05. Enrichment analysis for Gene Ontology terms. **(F)** Volcano plot displaying DEGs in WA treatment compared with control. Upregulated genes are colored in red, downregulated genes are colored in green, insignificantly altered genes are colored in blue. **(G)** Venn diagram of overlapping significantly changed genes (± 1.2 fold, P < 0.05). **(H)** The top ten overlapping genes are presented. **(I)** Protein–protein interaction network by GeneMANIA. **(J)** Heatmap of the top 120 significantly changed genes of subcutaneous adipose tissue from human under the condition of WAT browning induction (using GSE116801 dataset).

### Prdm16 and FATP1 mediate the WA-promoted WAT browning and weight loss

To evaluate the roles of Prdm16 and FATP1 in the WA-induced WAT browning and weight loss, the in vivo RNA interference (RNAi) technology was employed (**Figure 5A**). Prdm16 or FATP1 knockdown restored the adipocytic sizes and the expression levels of the browning-associated genes (**Figure 5B-D**), and also diminished the reduction in both body weight and fat pad weights (**Figure 5E-F**) in the WA-treated mice. The food intake was not affected by Prdm16 or FATP1 knockdown (**Figure 5G**). Knockdown of Prdm16 or FATP1 dampened both increased EE and decreased RER induced by WA (**Figure 5H-I; Figure S6A**). Intriguingly, we observed that Prdm16 knockdown remarkably downregulated FATP1 expression with or without WA treatment (**Figure 5J-K**). In contrast, FATP1 knockdown did not change Prdm16 expression level (**Figure 5J-K**). Collectively, these findings suggest that FATP1 may be a downstream effector of Prdm16, and this Prdm16-FATP1 axis may mediate the WAT browning-inducing and body weight-reducing effects of WA.

**Figure 5.**
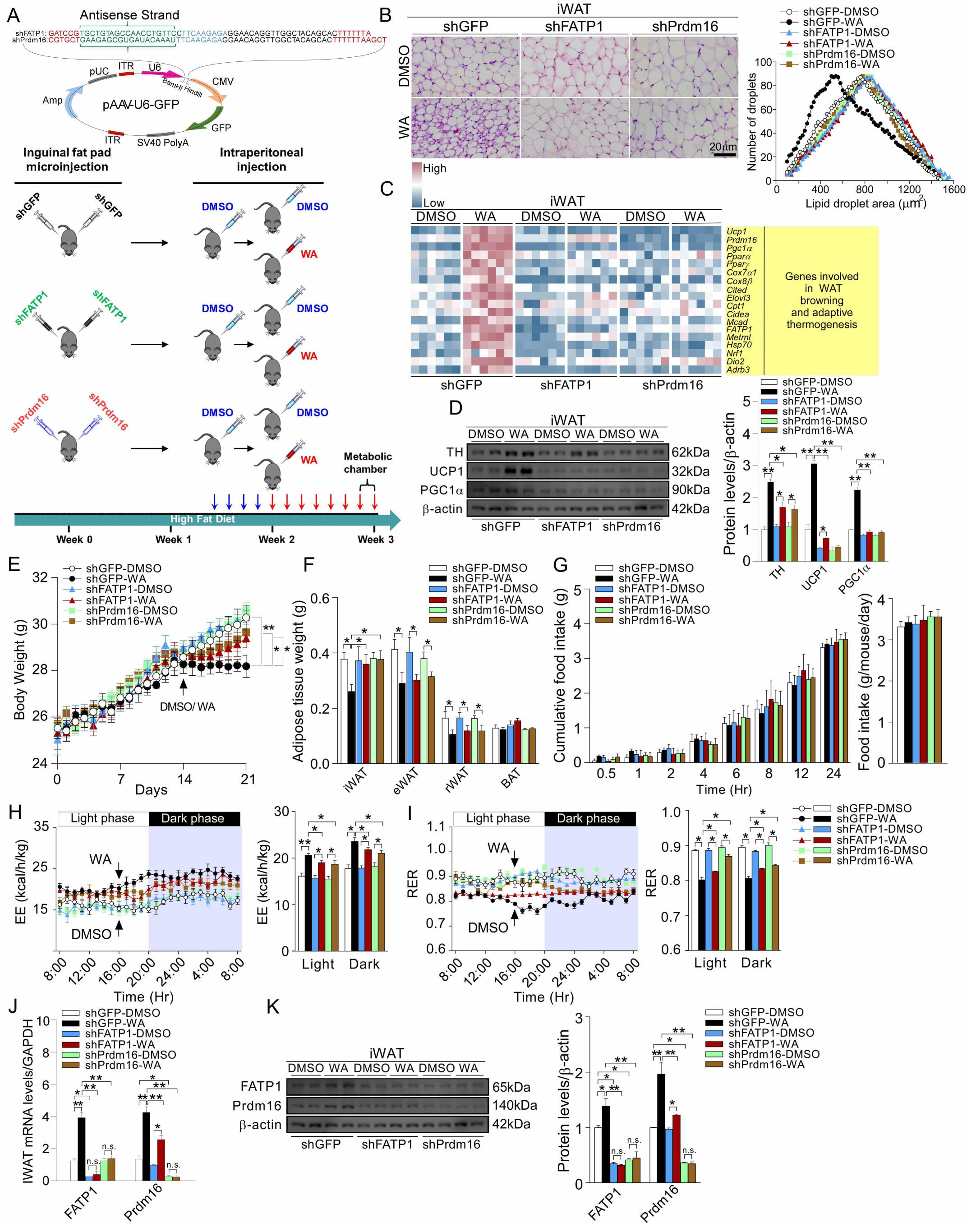
Prdm16 and FATP1 mediate the WA-promoted WAT browning and weight loss. 8 week-old mice were injected with shGFP, shPrdm16 or shFATP1, and then treated with WA or DMSO daily for 7 days (shGFP-DMSO, n = 7; shGFP-WA, n = 6; shFATP1-DMSO, n = 7; shFATP1-WA, n = 6; shPrdm16-DMSO, n = 11; shPrdm16-WA, n = 11). **(A)** Schematic illustration of experiments. Mice were injected with shGFP, shPrdm16 or shFATP1. 2 weeks later, mice were treated with WA or DMSO daily for 7 days. Tissue harvest was performed at week 3. **(B)** Left panel, H&E staining. Scale bar indicates 20 μm. Right panel, cell size profiling of adipocytes from iWAT. **(C)** Heatmap visualization of relative mRNA expression of the genes associated with WAT browning in iWAT (n = 6). **(D)** Western blot with quantification of TH, Ucp1 and Pgc1α in iWAT. **(E)** Body weight. **(F)** Adipose tissue weight. **(G)** Cumulative food intake. **(H-I)** Indirect calorimetry was performed to quantify EE (**H**) and RER (**I**) of shGFP, shPrdm16 or sh**FATP1**-injected mice treated with WA or DMSO during complete 24 hr light-dark cycles, the arrow indicates the time of WA or DMSO injection (n = 5). **(J)** Relative mRNA expression of TH, Ucp1 and Pgc1α in iWAT. **(K)** Western blot with quantification of FATP1 and Prdm16 in iWAT. Values were expressed as mean ± SEM. Significance was determined by one-way ANOVA with Tukey post hoc test. *p < 0.05, **p < 0.01, and ***p < 0.001.

### WA promotes WAT browning through sympathetic nerve-mediated activation of ***Prdm16-FATP1 axis***

To determine whether Prdm16-FATP1 axis may also be associated with the sympathetic activity in human adipocytes, we performed a correlation analysis between Prdm16, FATP1, PGC1α, and DβH (the enzyme converting dopamine to norepinephrine) using the GTEXv5 human subcutaneous and visceral adipose tissue databases. We observed that Prdm16 and FATP1 mRNA levels were positively correlated with both PGC1α and DβH (**Figure S7A-S7H**). This analysis verified that Prdm16 and FATP1 are also correlative with the sympathetic nervous activity and mitochondrial function in the human adipocytes. Collectively, these findings point to a model illustrated in **Figure 6**.

**Figure 6.**
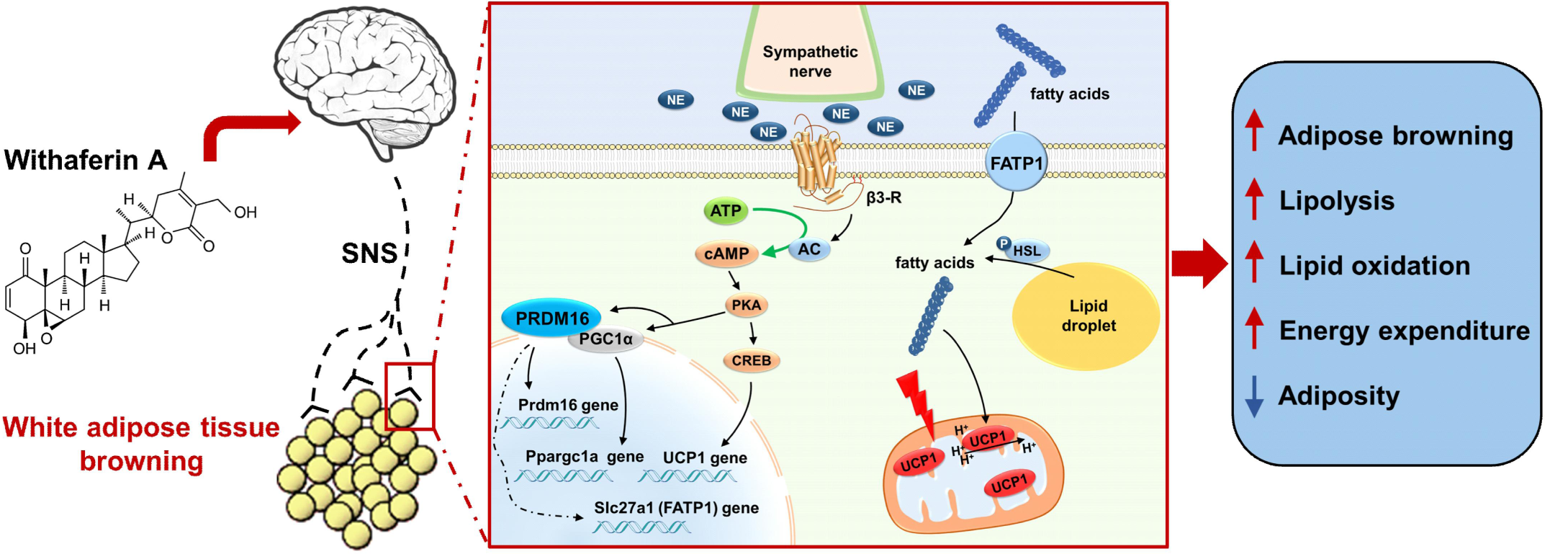
WA promotes WAT browning through sympathetic nerve-mediated activation of Prdm16-FATP1 axis. Schematic summary of the mechanism underlying the WAT browning-inducing and anti-obesity effects of WA: the WAT browning induced by WA is dependent on the activation of Prdm16 and FATP1 mediated by sympathetic nerves-WAT axis.

In conclusion, this study demonstrates the therapeutic potential of WA for obesity treatment, and reveals that Prdm16 may be the main transcription factor responsible for WA-induced adipose browning through sympathetic nerve-adiposity axis; and FATP1 may be the main fatty acid transporter in downstream of Prdm16. These findings indicate that WA promotes WAT browning and prevents HFD-induced obesity through sympathetic nerve-mediated activation of adipocytic Prdm16-FATP1 pathway.

## Discussion

WA is a liposoluble natural compound with an ability to cross the BBB. WA exerts profound neuropharmacological effects, including promotion of neurite outgrowth, mitigation of neuritic atrophy, and facilitation of synapse reconstruction (Dar et al., 2015; Zahiruddin et al., 2020). As a newly-recognized leptin sensitizer, WA prevents obesity by potentiating hypothalamic leptin signaling (Lee et al., 2016). However, the mechanism by which WA prevents obesity remains to be elucidated.

In this study, we found that the intraperitoneal treatment of WA promoted WAT browning, elevated EE and prevented high fat diet-induced obesity, without affecting food intake. Our long-term study showed that WA exerted sustained anti-obesity effects over a period of two months. Emerging evidence indicates that WA exerts its unique anti-obesity effects by enhancing the hypothalamic leptin signaling. WA (1.25 mg/kg, i.p. once daily) leads to an increased leptin sensitivity and a reduced body weight in DIO mice, but not in lean mice, leptin-deficient (ob/ob) mice or leptin receptor-deficient (db/db) mice (Lee et al., 2016). WA (1.25 mg/kg, i.p. once daily) increases the level of p-STAT3 (Tyr705), reduces the level of p-PERK (Thr980) in hypothalamus, demonstrating that WA relieves ER stress, sensitizes leptin signaling (Lee et al., 2016). In this study, we also observed that WA upregulated the expression levels of the genes associated with leptin signaling in the hypothalamus, confirming that the enhancement of leptin sensitivity indeed contributes to the anti-obesity effects of WA.

Multiple lines of evidence show that the hypothalamus-SNS axis plays crucial roles in the regulation of WAT browning (Jeremic et al., 2017; Kwon et al., 2016). A series of anti-obesity hormones (leptin, adiponectin), pharmacological agents (butein, phyllodulcin), environmental stimuli (cold exposure), and physical exercise promote WAT browning through hypothalamus-SNS axis (Dodd et al., 2015; Kim et al., 2017a; Shin et al., 2017; Song et al., 2018). The visualization of sympathetic neuro-adipose connections strengthens the notion that sympathetic nerves are essential for beige adipogenesis (Hu et al., 2020; Zeng et al., 2015; Zeng et al., 2019). We found that WA elevated EE by 10.1% in the bilateral sympathetic denervated-mice, whereas WA elevated EE by 25.3% in sham-operated mice. These observations suggest that WA-induced WAT browning by sympathetic nerve-adiposity connection essentially contributes to the anti-obesity effects of WA.

The sympathetic nerves regulate the adipocytic factors associated with WAT browning. Stimulation of sympathetic nerves increases the level of connexin 43 in WAT (Kim et al., 2017b); and WAT specific connexin-43 overexpression promotes WAT browning (Zhu et al., 2016). In this study, we observed that Prdm16 and FATP1 in iWAT were upregulated by WA, and this was blocked by sympathetic denervation. The inhibition of Prdm16 or FATP1 expressions in iWAT diminished the WAT browning-inducing and body weight-reducing effects of WA. Moreover, the gene expression profile analysis of human subcutaneous fat (Haidich, 2010) also revealed that Prdm16 and FATP1 are critical factors for the maintenance of fat homeostasis in human.

Prdm16 is a key adipocytic transcription factor governing the development of beige adipocytes (Cohen et al., 2014; Inagaki et al., 2016; Seale et al., 2008; Seale et al., 2007). Stimulation of sympathetic nerves activates Prdm16 in white adipocytes via β3R signaling, which further activates Pgc1α/Pparα/Pparγ/retinoid X receptor pathway to upregulate UCP1, leading to WAT browning (Iida et al., 2015; Inagaki et al., 2016). The fatty acid transporters (FATPs) in white adipocytes are activated in response to the conditions that induce WAT browning. FATPs are the main fatty acid suppliers in beige adipocytes (Anderson and Stahl, 2013; Calderon-Dominguez et al., 2016; Henkin et al., 2012). The activation of FATP1 enhances WAT browning and protects against HFD-induced obesity (Cao et al., 2019; Xu et al., 2017). In human, the polymorphisms and mutations in FATP genes (FATP1, FATP4 and FATP5) are associated with the pathological lipid metabolism and insulin signaling, showing that FATPs are potential drug targets for obesity and T2DM therapies (Auinger et al., 2010; Gertow et al., 2004; Meirhaeghe et al., 2000; Rask-Andersen et al., 2011). These findings suggest that Prdm16-FATP1 axis is essential for mediating WAT browning induced by WA.

Mounting evidence indicates that the external stimuli that promote WAT browning may also lead to an alteration in fatty acid composition in adipose tissues (Srivastava and Veech, 2019). Importantly, the modification of the fatty acid composition in WATs contributes to the promotion of WAT browning, which has been considered a novel strategy for obesity treatment (Chaurasia et al., 2016; Shen et al., 2013); (Li et al., 2018). The unsaturated fatty acids, such as oleic acid and arachidonic acid, facilitate the mitochondrial uncoupling and oxidative phosphorylation in adipocytes, contributing to the prevention of obesity, fatty liver disease and cardiovascular disease (Croset et al., 2000; Kantae et al., 2017; Matsuda et al., 2020; Vögler et al., 2008). We found that WA increased the levels of oleic acid and arachidonic acid, and reduced the level of tricosanoic acid in iWAT, and these were normalized by sympathetic denervation. Therefore, we can not rule out that the modification of fatty acid composition caused by WA may also be implicated in the anti-obesity or the metabolic beneficial effects of WA.

In summary, our findings demonstrate that WA prevents HFD-induced obesity by promoting WAT browning through sympathetic-adipose axis. Prdm16-FATP1 pathway in adipocytes mediates the WA-induced WAT browning. Our results suggest that WA could be a candidate of anti-obesity agents, and provide a new insight into obesity treatment and the maintenance of human health.

## Materials and methods

### Mice

Mice were housed under controlled light (12-h dark-light cycle, with the dark cycle encompassing 8 p.m. to 8 a.m), temperature (22 ± 2 °C) conditions with a free access to food and water. For diet-induced obesity studies, mice were were placed on 60 kcal% High Fat Diet (HFD, Research Diets, D12492i) at the age of 8 weeks. For thermoneutral studies, mice were housed at 30 °C in a light-controlled climatic chamber. During all procedures of experiments, the number of animals and their suffering by treatments were minimized.

### Reagents

Ucp-1, Pgc1α, TH, FATP1, Prdm16 and β-actin specific antibodies were purchased from Abcam (ab-10983, San Francisco, CA), Santa Cruz Biotechnologies (sc-13067, Santa Cruz, CA), Millipore (AB152, Billerica, MA), Signalway Antibody (45112, College Park, MD), Signalway Antibody (25030, College Park, MD), Sigma (A5316, St Louis, MO). Anti-rabbit IgG / Alexa Fluor 488 were purchased from Bioss (bs-02950-AF488). Tranzol up, TransScript one-step gDNA removel and cDNA synthesis Super MiX, TransStart Top Green qPCR Super Mix were purchased from TransGen Biotech. RNaseZapTM were purphased from Invitrogen. Triglycerides (TG), Cholesterol (CHO), High density lipoprotein cholesterol (HDL-C) and Low density lipoprotein cholesterol (LDL-C) Regen Kit were purchased from BioSINO Bio-thchnology. ECL assay kit were purchased from Bio-Rad (California, CA). WA were purchased from ChromaDex (Irvine, CA). Dimethyl sulfoxide (DMSO) were purphased from Amresco (Solon, CA). The neurotoxin (6-hydroxydopamine) was purchased from Sigma-Aldrich (6-OHDA, H4381, St. Louis, MO).

### Administration of WA

Administration of WA was carried out as previously described (Lee et al., 2016). For intraperitoneal (i.p.) treatment, mice received 25 μl of vehicle (DMSO) for four days as acclimation before WA treatment. Then, WA was dissolved in DMSO (25 μl; 0.2, 2, 20, 200 μg/kg) and administered to mice once a day for 7, 14, 21 days or 2 months. Vehicle groups received 25 μl of DMSO during the course of experiments. All treatments were performed within 90 min before dark cycle.

### Food intake and body weight measurements

Food intake and body weight were measured daily, and the percent increase of body weight was calculated by the following equation: 100 X (body weight of WA or vehicle injected group - initial body weight) / (initial body weight).

### Infrared thermal imaging

Infrared thermal imaging (IRT) was performed using a FOTRIC thermal camera. Each mouse was placed on a cage top at a fixed distance away from the camera lens. Serial 1-s images (10 Hz) were taken in triplicate at baseline and at 15-min intervals for 1 h after intraperitoneal injection of WA or vehicle.

IRT images were analyzed using AnalyzIR software. For analysis, a constantly sized circular region of interest was drawn over iWAT and BAT, and the average temperature was recorded.

### Indirect calorimetry

After an adaptation to single caging, miced were placed in metabolic chambers on the day 6 of WA treatment. During and after the 24 h acclimation, we administered vehicle or WA (2 μg/kg) within 90 min before dark cycle. Indirect calorimetry recording was performed using an indirect open-circuit calorimeter Oxylet Physiocage System (LE1305 Physiocage 00; LE405 O2/CO2 Analyzer; LE400 Air Supply and Swithching; Panlab, Cornella, Spain). Room air flowed through each chamber at a rate of 450 ml/min. The O2 and CO2 levels were measured during 3 min sampling periods every 30 min and data were analyzed with the METABOLISM software (v2.2.01). Locomotor activity were measured using a 2-dimensional infrared light-beam. The VO2 and VCO2 were expressed in ml/min/kg. The RER was determined by the ratio VCO2/VO2. The EE was calculated according to following equation: EE (kcal/day/kg) = VO2 × 1.44 × [3.815 + (1.232 × RER)]. The mean values for VO2, VCO2, RER and EE of dark cycle and light cycle were compared for each group.

### Sympathetic denervation

Sympathetic denervation was carried out as previously described (Dodd et al., 2015). 8-week-old male mice received 20 microinjections of 6-hydroxydopamine [6-OHDA; 1 μl per injection, 9 mg/ml in 0.15 M NaCl containing 1% (w/v) ascorbic acid] throughout the right or both inguinal fat pads. Sham operated fat pads received an equal volume of vehicle. Two weeks (unilateral) or 5 weeks (bilateral) after 6-OHDA injections, mice were i.p. injected with WA (2 μg/kg) once a day for 7 days. Body weights were monitored throughout the duration of the experiment. Inguinal WATs were harvested for histological/immunofluorescence assessment, or processed for western analysis, real-time qPCR and whole-genome sequencing analysis.

### Total protein extraction and western analysis

Inguinal WATs were homogenized with a Polytron in ice-cold RIPA buffer (1% Trinton X 100; 10 mM Na2HPO4; 150 mM NaCl; 1% DOC; 5 mM EDTA; 5 mM NaF; 0.1 % SDS) supplemented with protease and phosphatase inhibitors (catalog #P8340 and #P2850; Sigma), sonicated and cleared by centrifugation (10,000 × g, 10 min, at 4 °C). Protein concentration in the supernatant was determined by BCA assay (Aidlab; PP01). Protein (5 μg) in 1 × sample buffer [62.5 mM Tris·Cl (pH 6.8), 2% (wt/vol) SDS, 5% glycerol, 0.05% (wt/vol) bromophenol blue] was denatured by boiling at 100°C for 5 min and separated on 8% sodium dodecyl sulfate poly acrylamide (SDS-PAGE) gels and transferred onto nitrocellulose membrane (Pall Corporation; T60327) by electrophoresis. After electrophoresis, blots were blocked in 5% nonfat milk in Tris-buffered saline and Tween 20 (TBST) for 2 h at room temperature and probed with primary antibody in 5% BSA-TBST overnight at 4°C. After primary incubation, the blots were washed three times in TBST for 15 min, followed by incubation with HRP-conjugated secondary antibody in TBST with 5% nonfat milk for 2 h at room temperature. Then, the membranes were washed three times for 15 min with TBST, and developed the membranes using an Enhanced Chemiluminescence assay (BIO-Rad) and quantified band intensities using ImageJ software (NIH).

### Quantitative real-time PCR

Total RNA for quantitative real-time PCR (qPCR) was extracted from fat pads and hypothalamus using TRIzol reagent (TransGen Biotech) by following the manufacturer’s protocol. Quantification and integrity analysis of total RNA was performed by running 1 µl of each sample on NanoDrop 5500 (Thermo). Complementary DNA (cDNA) was prepared by reverse transcription (TransScript one-step gDNA removel and cDNA synthesis Super MiX, TransGen Biotech). The relative expression of mRNAs was determined by the SYBR Green PCR system (Bio-Rad). The relative expression of genes of interest was calculated by comparative Ct method and GAPDH was used as an endogenous control. Sequences of the primers used for real-time qPCR are available in Supplementary table 1.

### Hematoxylin and eosin (H&E) staining

Inguinal WATs were collected from mice treated with WA (2 μg/kg) or vehicle for 7 days and immediately fixed in 4% paraformaldehyde solution for 48 h, then the samples were embedded in optimal cutting temperature OCT compound (Sakura FineTech, Tokyo). Tissue sections of 10 μm thickness were stained with hematoxylin and eosin (H&E). Stained slides were analyzed at the indicated magnification, and images were captured by a digital camera (Olympus).

### Measurement of adipocyte size

Inguinal WAT sections were stained with H&E. Three mice per group were randomly selected. Adipocyte images were acquired using a microscope (Olympus) at 200 × magnification. The cellular size was quantified by using Image J software, and then the number of adipocytes falling into each field with intervals of 5 μm was counted.

### Immunohistofluorescence staining

Inguinal WATs were collected from mice treated with WA (2 μg/kg) or vehicle for 7 days and immediately fixed in 4% paraformaldehyde solution for 48 h. Then, the samples were incubated sequentially with 20% sucrose and 30% sucrose in PBS for 2 d and frozen in OCT compound (Sakura FineTech, Tokyo). Tissue sections of 10 μm thickness were taken on a cryostat and allowed to air dry on slides, followed by processing or preservation at -80 °C according to standard procedure. Frozen sections of tissues were subjected to TH staining. Sections were washed in PBS for 10 min, followed by incubation in blocking solution (10% normal goat serum; 0.2% Triton X-100; 2% BSA; PBS) for at least 1 h at room temperature. Primary antibodies [antibody anti-TH (1:200)] were applied in blocking solution and incubated overnight at 4 °C. Sections were washed at least three times with 5-min incubations in PBS plus 0.2% Triton X-100. Then a Alexa-Fluor 488-conjugated secondary antibody (1:200) was applied in blocking solution and incubated at room temperature for 2 h, followed by five washes with PBS plus 0.2% Triton X-100; and nuclei were stained with 4’,6-diamidino-2-phenylindole (DAPI). Sections were mounted with VectaShield medium (Vector Laboratories) and analyzed on a microscope.

### Whole-genome sequencing analysis

Total RNA was extracted from inguinal WATs of WA or vehicle-treated mice using TRIzol reagent (TransGen Biotech). The quality of the RNA was determined with NanoDrop 5500 (Thermo). For library preparation, 3 µg of total RNA per sample was used. Sequencing libraries were generated with NEBNext® Ultra™ RNA Library Prep Kit for Illumina® (NEB, USA). RNA molecules were selected using poly-T oligo attached magnetic beads, fragmented, and reverse transcribed with the Elute, Prime and Fragment Mix. Then, end repair, A-tailing, adaptor ligation, and library enrichment were performed according to the manufacturer’s instructions. RNA libraries were assessed for quality using the Agilent 2100 BioAnalyzer. The clustering of the index-coded samples was performed on a cBot Cluster Generation System using TruSeq PE Cluster Kit v3-cBot-HS (Illumia). Then, the RNA libraries were sequenced as 100 bp/50 bp paired-end runs on an Illumina Hiseq 2000/2500 platform. Differential expression analysis of two conditions/groups (two biological replicates per condition) was performed using the DESeq R package (1.10.1). The resulting P-values were adjusted using the Benjamini and Hochberg’s approach for controlling the false discovery rate. Genes with an adjusted P-value < 0.05 found by DESeq were assigned as differentially expressed. Gene Ontology (GO) enrichment analysis of differentially expressed genes was performed using the GOseq R package, in which gene length bias was corrected. GO terms with corrected P-value less than 0.05 were considered significantly enriched by differential expressed genes. Samples were measured and analyzed in Novogene Bioinformatics Technology Co., Ltd.

### AAV9-GFP construction and vectors injection

Prdm16 knockdown, FATP1 knockdown and control recombinant Adeno-associated virus-green fluorescence protein (AAV-GFP) vectors were constructed (Vigene Biosciences). Vectors injection procedures were performed as previously described (Zhang et al., 2017). Briefly, 8-week-old male mice were injected at multiple sites of the inguinal WAT pads (Virus diluted in sterile PBS: 1 × 10^10^ PFU/100 μl). Two weeks after vectors injections, mice were i.p. injected with WA (2 μg/kg) once a day for 7 days. Body weights were monitored throughout the duration of the experiment. Inguinal WATs were harvested for histological/ immunofluorescence assessment, or processed for western analysis, and real-time qPCR.

### Blood collection

Whole blood was collected by cardiac puncture and transferred to ice-cold EP tubes. The tubes were centrifuged at 2000 x g for 30 min at 4°C and stored at -80°C. The serum was used for lipid measurement.

### Lipidomics analysis

For lipidomic analysis, the extraction of lipids was carried out using the liquid–liquid extraction protocol. Briefly, 50 mg frozen iWAT tissues was dissected followed by addition of 2.5 ml of mixture of chloroform and methanol (V:V = 2:1) on ice, and the mixture was homogenized followed by addition of 1.25 ml distilled water and vortex for 30 s and then incubated on ice for 5 min. Then, the sample was centrifuged at 12,000 x g at 4 °C for 10 min. The lower chloroform layer was collected and dried by nitrogen. Samples were reconstituted in chloroform and methanol solution (V:V = 2:1), separated with Cortecs C18 column (2.1 × 100 mm, Waters) and analyzed by an Q Exactive orbitrap mass spectrometer that was operated in both positive and negative mode using Xcalibur 3.0 softare. A full-scan followed by 10 data-dependent MS/MS scans were acquired using higher energy collisional dissociation with stepped normalized collision energy of 15%, 30% and 45%. The data analysis was performed on LipidSearch softare (Thrmo, Waltham, MA).

### Bioinformatic analysis of human dataset

A subcutaneous adipose tissue transcriptomic dataset associated with WAT browning and obesity was identified (GSE116801). Log2-transformation was applied as needed. Heatmap of the top 120 significantly changed genes of subcutaneous adipose tissue from human under the condition of WAT browning induction was carried out using the R/GeneMeta package. ***Statistical analysis.*** All data are expressed as the Mean ± SEM. Statistical significance was determined using Student’s t test (two-tailed) or one-way or two-way ANOVA, as indicated in the figure legends. Numbers of cohorts and n values for each experiment are indicated in the figure legends. All data were analyzed using the appropriate statistical analysis methods with the SPSS software (version 19.0). The analysis of EE, VO2 and VCO2 in Supplementary figure 2, 3, 4 were based on ANCOVA. Significance was accepted at * P < 0.05, ** P < 0.01 or *** P < 0.001.

### Data availability

The data set generated in this study is available from the corresponding author on reasonable request. Accession codes will be available before publication.

## Conflict of interest

The authors declare that they have no conflict of interest.

## Author Contributions

B.G. and J.L. did the experiment, analyzed the data. B.W., C.Z., Z.S., and M.Z. participated in experiments. J.L. and R.Z. wrote the manuscript. R.Z. conceived the idea of study and manuscript writing. All authors reviewed and approved the manuscript for submission.

---

**Supplementary Figure 1.**
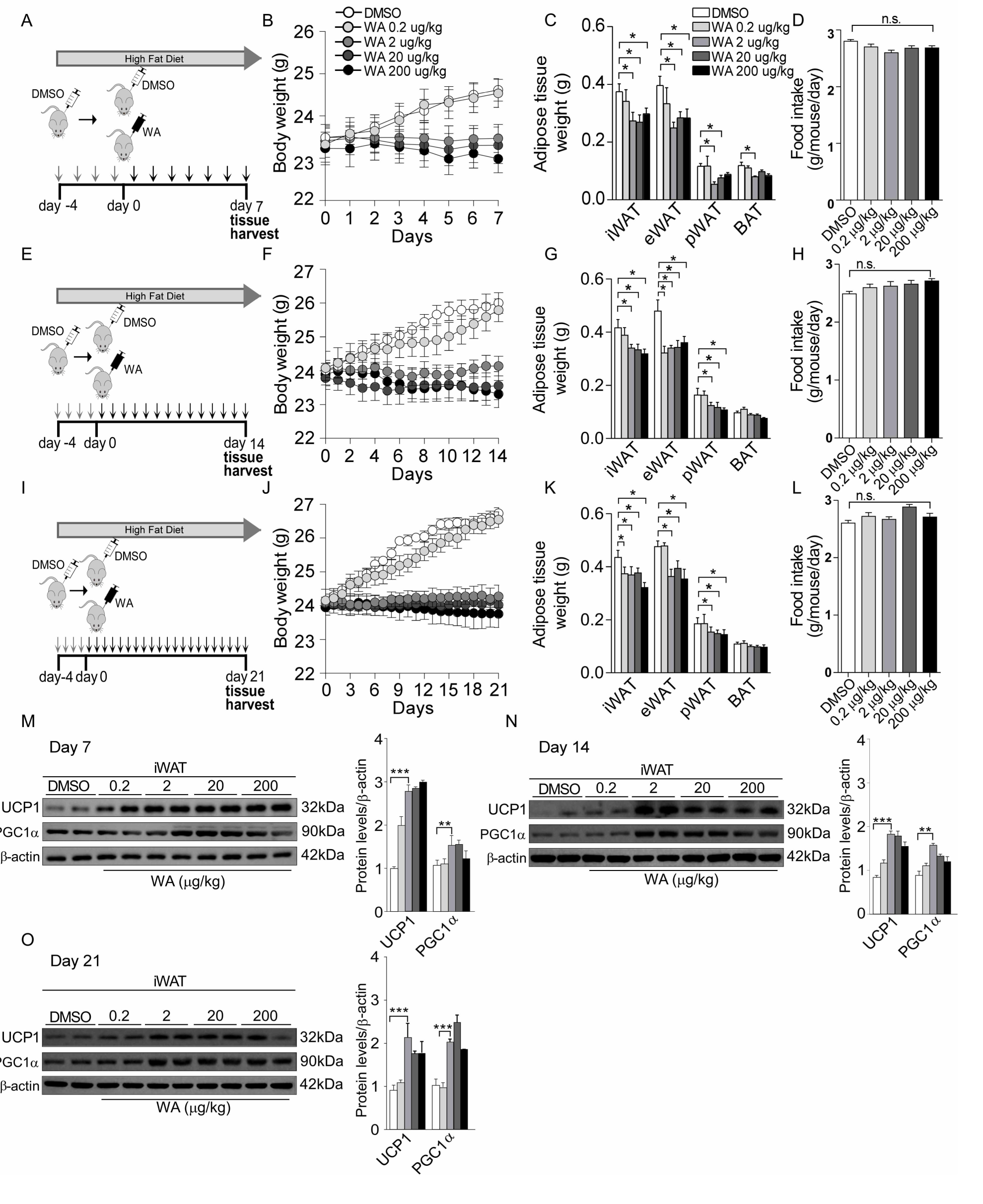
A 3-week dose-ranging study of WA. **(A-D)** Mice were housed at 22°C, fed a HFD, and treated with WA at doses ranging from 0 to 200 μg/kg for 7 days (DMSO, n = 10; WA 0.2 μg/kg, n = 5; WA 2 μg/kg, n = 10; WA 20 μg/kg, n = 8; WA 200 μg/kg, n = 5). **(A)** Schematic illustration of experiments. Mice received DMSO for four days as acclimation, then they were treated with WA or DMSO for 7 consecutive days. **(B)** Body weight. **(C)** Adipose tissue weight. **(D)** Food intake. **(E-H)** Mice were housed at 22°C, fed a HFD, and treated with WA at doses ranging from 0 to 200 μg/kg for 14 days (DMSO, n = 9; WA 0.2 μg/kg, n = 5; WA 2 μg/kg, n = 10; WA 20 μg/kg, n = 8; WA 200 μg/kg, n = 5). **(E)** Schematic illustration of experiments. Mice received DMSO for four days as acclimation, then they were treated with WA or DMSO for 14 consecutive days. **(F)** Body weight. **(G)** Adipose tissue weight. **(H)** Food intake. **(I-L)** Mice were housed at 22°C, fed a HFD, and treated with WA at doses ranging from 0 to 200 μg/kg for 21 days (DMSO, n = 10; WA 0.2 μg/kg, n = 5; WA 2 μg/kg, n = 9; WA 20 μg/kg, n = 9; WA 200 μg/kg, n = 5). **(I)** Schematic illustration of experiments. Mice received DMSO for four days as acclimation, then they were treated with WA or DMSO for 21 consecutive days. **(J)** Body weight. **(K)** Adipose tissue weight. **(L)** Food intake. **(M-O)** Western blot with quantification of Ucp-1 and PGC1α in iWAT of mice treated with WA for 7 days **(M)**, for 14 days **(N)**, and for 21 days **(O)**. Values are mean ± SEM. Significance was determined by one-way ANOVA with Dunnett multiple comparisons. *p < 0.05, **p < 0.01, and ***p < 0.001.

**Supplementary Figure 2 (related to figure 1).**
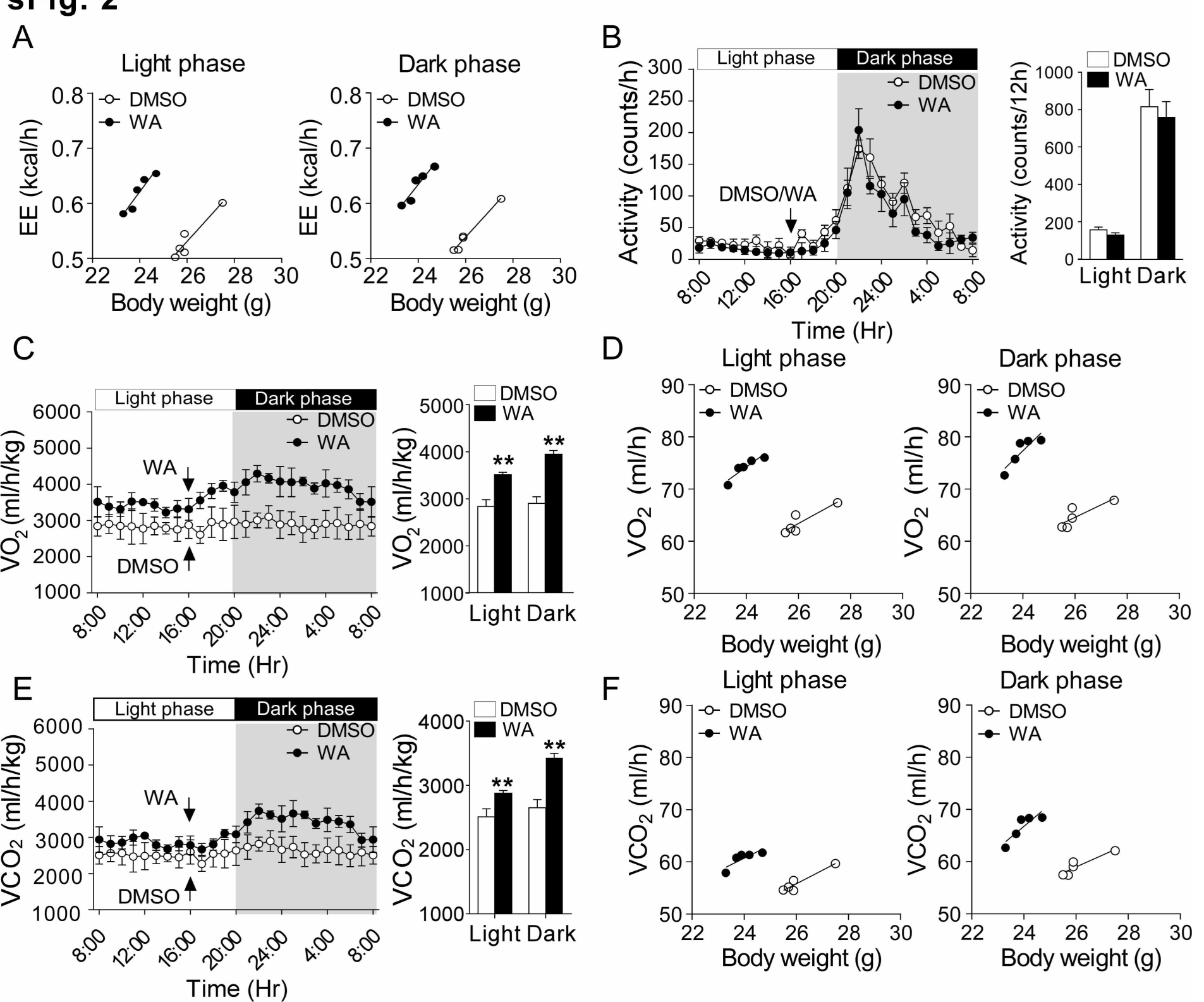
Analysis of VO2, VCO2 and EE in mice treated with WA. **(A)** EE of WA or DMSO-treated mice (n = 5). EE per whole animal is plotted against body weight. In all panels, lines show fitted regressions. **(B)** Indirect calorimetry was performed to quantify the motor activity of WA or DMSO-treated mice during complete 24 hr light-dark cycles, the arrow indicates the time of WA or DMSO injection (n = 5). **(C, D)** Oxygen consumption (VO2) (n = 5). VO2 per whole animal is plotted against body weight. In all panels, lines show fitted regressions. **(E, F)** Carbon dioxide production (VCO2) (n = 5). VCO2 per whole animal is plotted against body weight. In all panels, lines show fitted regressions. Values are mean ± SEM. Significance was determined by Student’s t test. **p < 0.01.

**Supplementary Figure 3 (related to figure 3).**
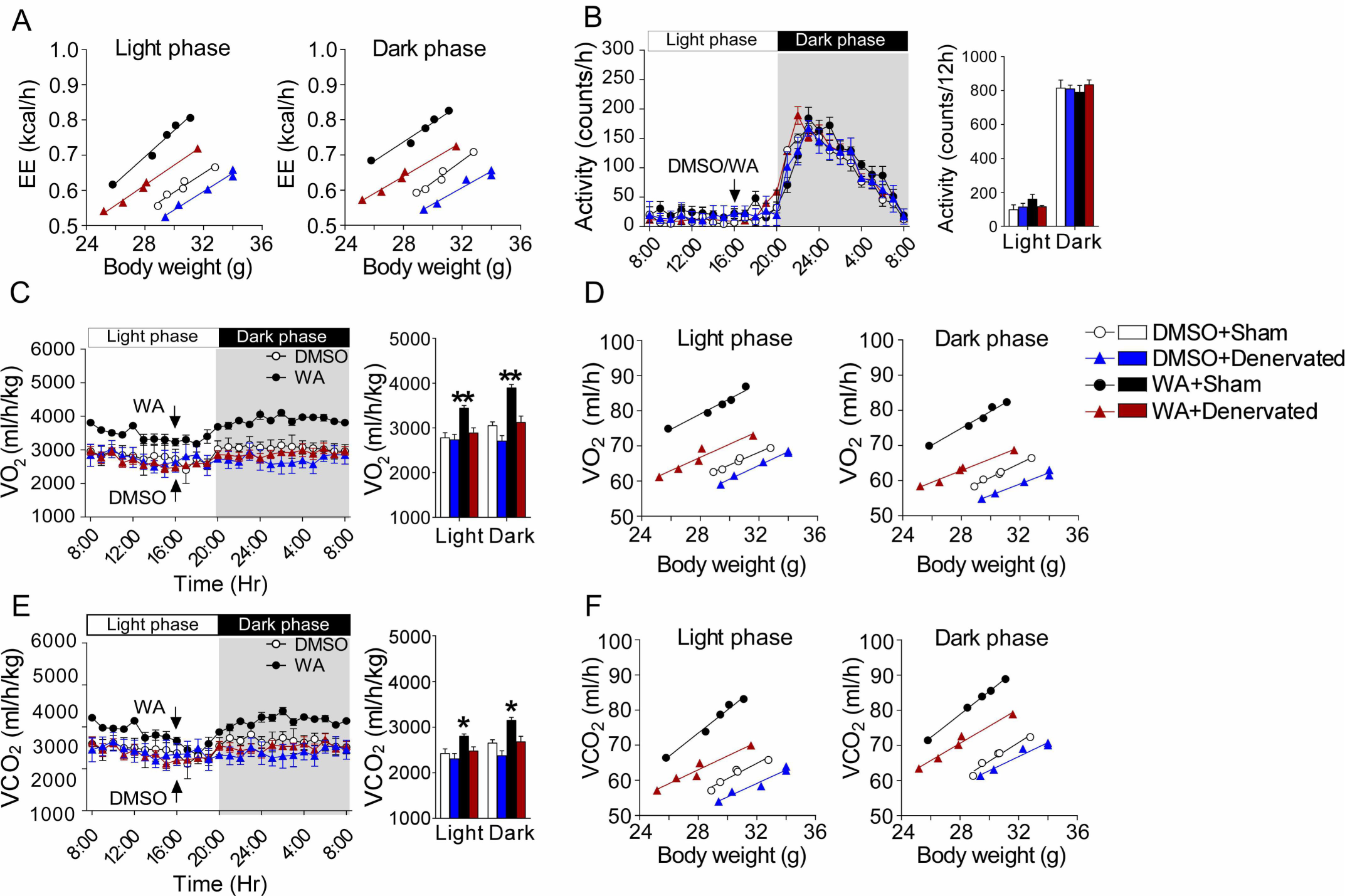
Analysis of VO2, VCO2 and EE in denervated or sham-operated mice treated with WA. **(A)** EE of denervated or sham-operated mice treated with WA or DMSO (n = 5). EE per whole animal is plotted against body weight. In all panels, lines show fitted regressions. **(B)** Indirect calorimetry was performed to quantify the motor activity of denervated or sham-operated mice treated with WA or DMSO during complete 24 hr light-dark cycles, the arrow indicates the time of WA or DMSO injection (n = 5). **(C, D)** VO2 of denervated or sham-operated mice treated with WA or DMSO (n = 5). VO2 per whole animal is plotted against body weight. In all panels, lines show fitted regressions. **(E, F)** VCO2 of denervated or sham-operated mice treated with WA or DMSO (n = 5). VCO2 per whole animal is plotted against body weight. In all panels, lines show fitted regressions. Values are mean ± SEM. Significance was determined by one-way ANOVA with Bonferroni test. *p < 0.05; **p < 0.01.

**Supplementary Figure 4 (related to figure 5).**
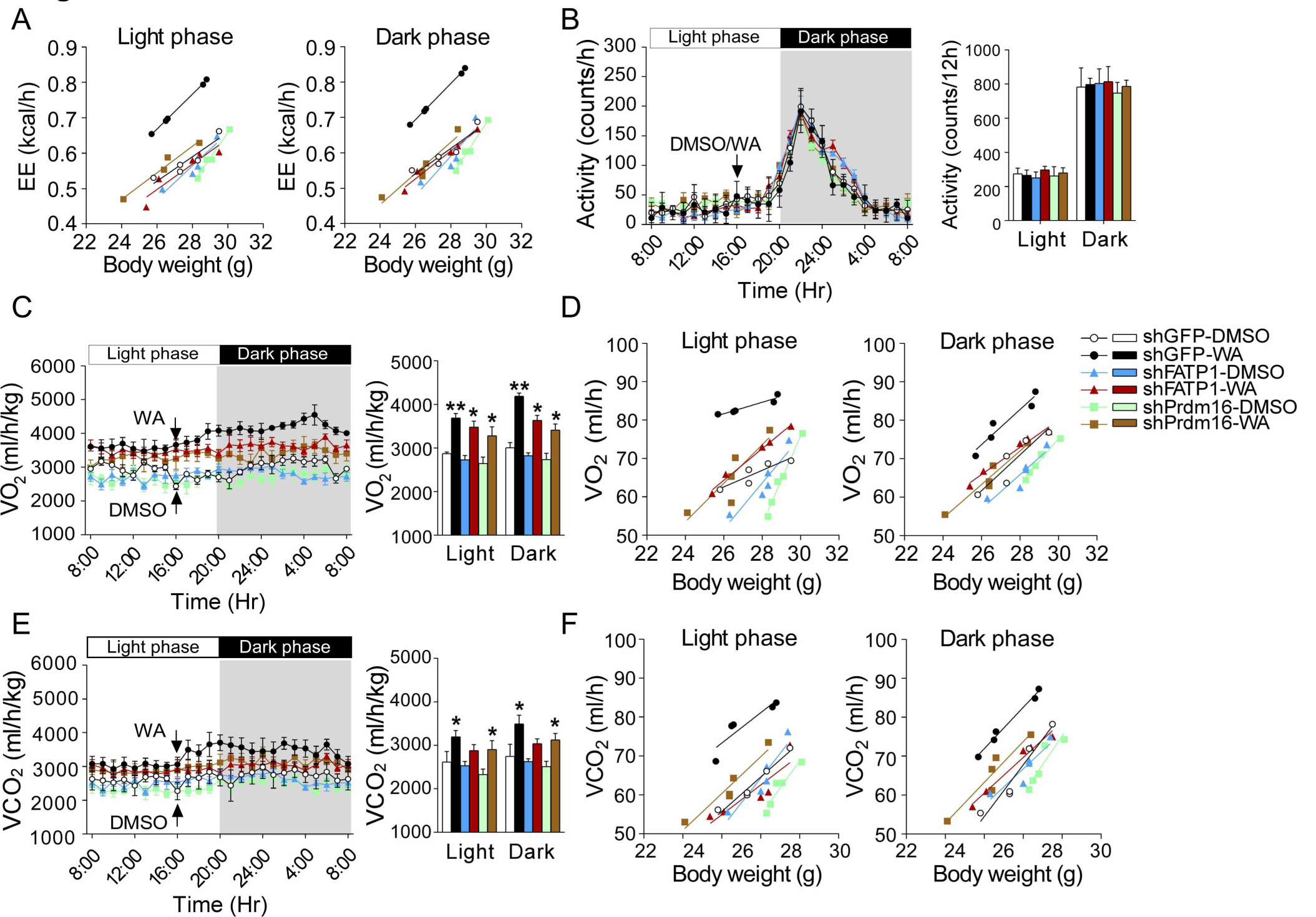
Analysis of VO2, VCO2 and EE in shGFP, shPrdm16 or shFATP1-injected mice treated with WA. **(A)** EE of shGFP, shPrdm16 or shFATP1-injected mice treated with WA or DMSO (n = 5). EE per whole animal is plotted against body weight. In all panels, lines show fitted regressions. **(B)** Indirect calorimetry was performed to quantify the motor activity of shGFP, shPrdm16 or shFATP1-injected mice treated with WA or DMSO during complete 24 hr light-dark cycles, the arrow indicates the time of WA or DMSO injection (n = 5). **(C, D)** VO2 of shGFP, shPrdm16 or shFATP1-injected mice treated with WA or DMSO (n = 5). VO2 per whole animal is plotted against body weight. In all panels, lines show fitted regressions. **(E, F)** VCO2 of shGFP, shPrdm16 or shFATP1-injected mice treated with WA or DMSO (n = 5). VCO2 per whole animal is plotted against body weight. In all panels, lines show fitted regressions. Values are mean ± SEM. Significance was determined by one-way ANOVA with Tukey post hoc test. *p < 0.05; **p < 0.01.

**Supplementary Figure 5.**
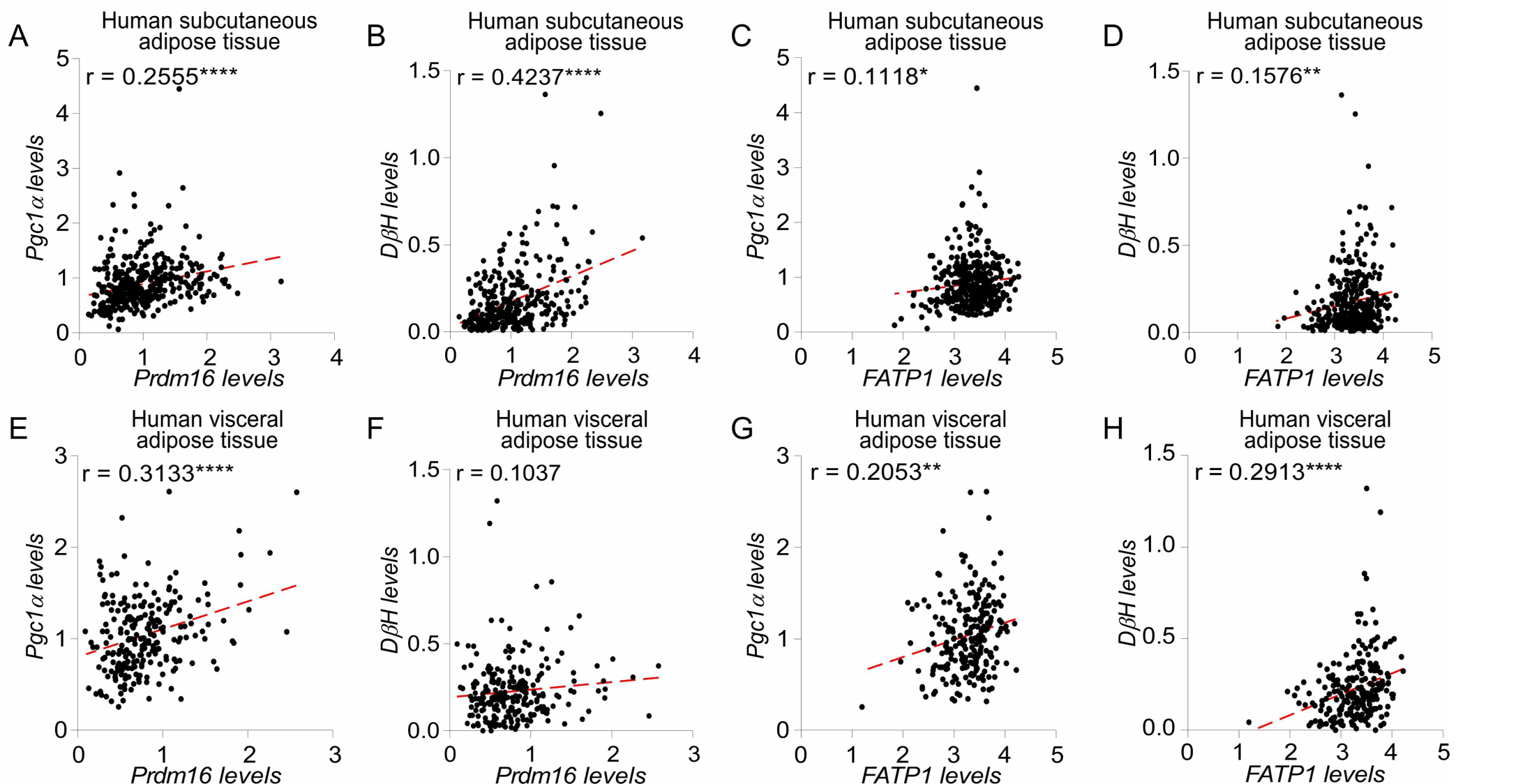
Correlation analysis between Prdm16, FATP1, Pgc1α and DβH in human adipose tissue. (Matsuda et al.) Correlation analysis between Prdm16, FATP1, Pgc1α and DβH in human subcutaneous (**A-D**) and visceral (**E-H**) adipose tissues (using GTEXv5 databases).

**Supplementary Figure 6.**
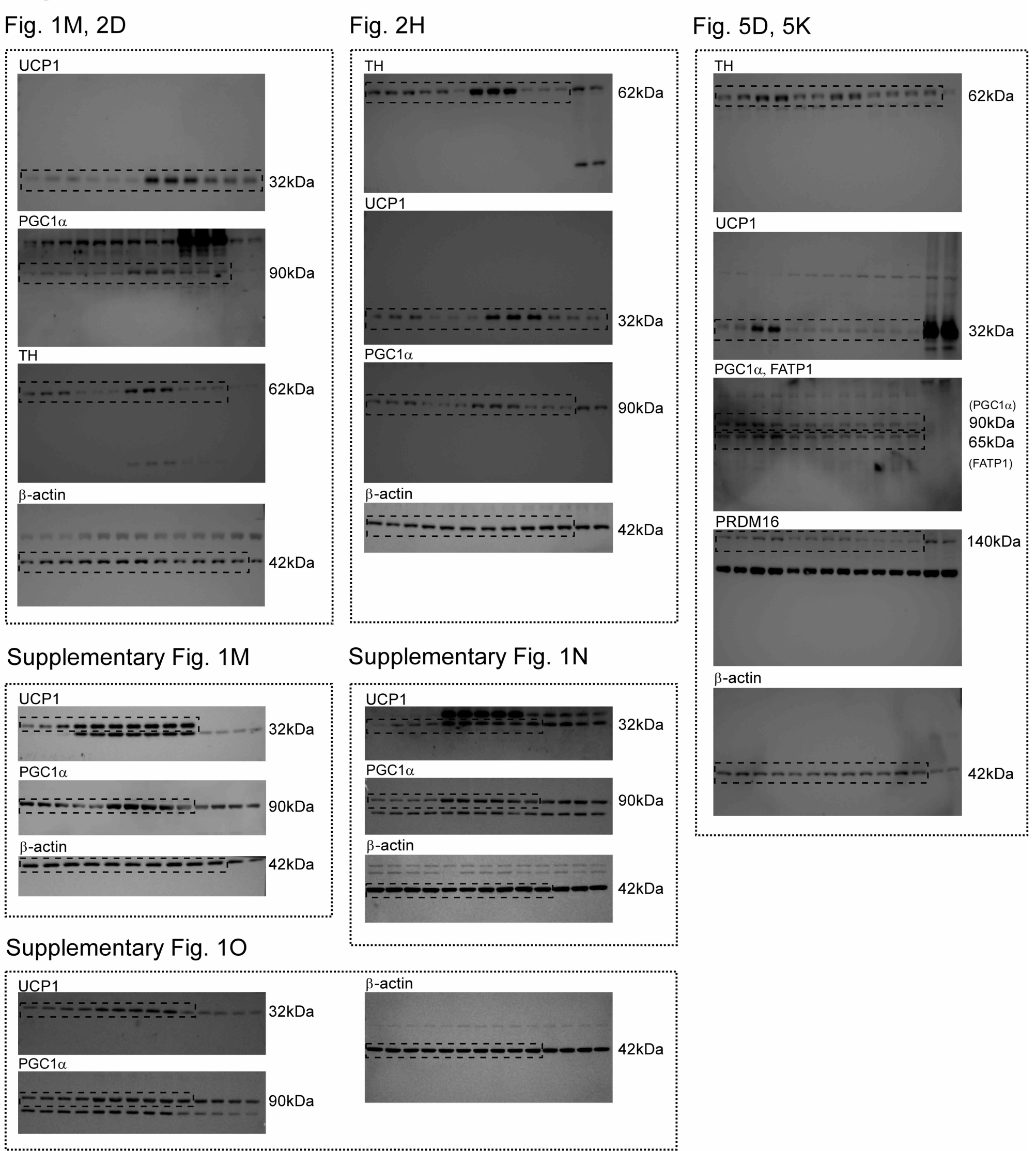
Original full western blot images

**Supplementary table 1:**
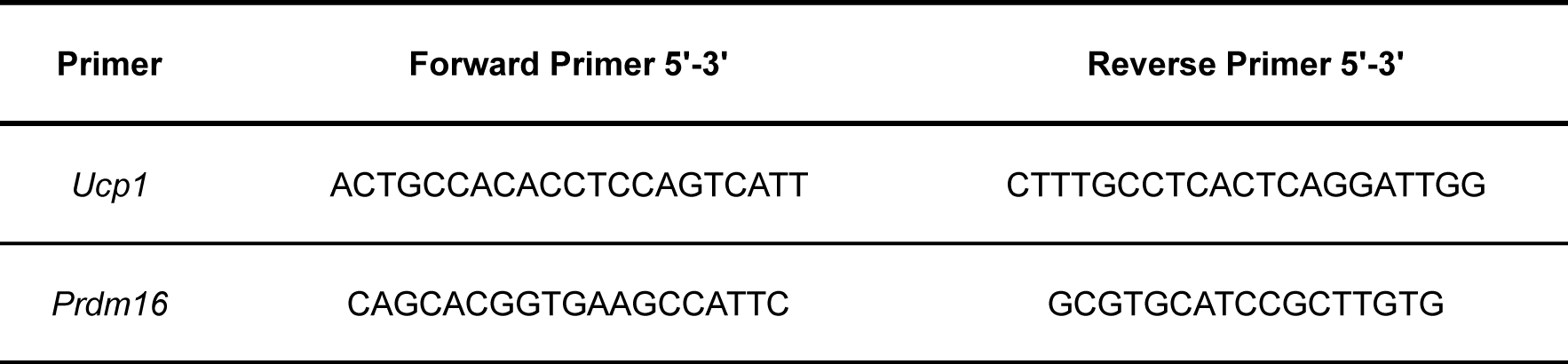
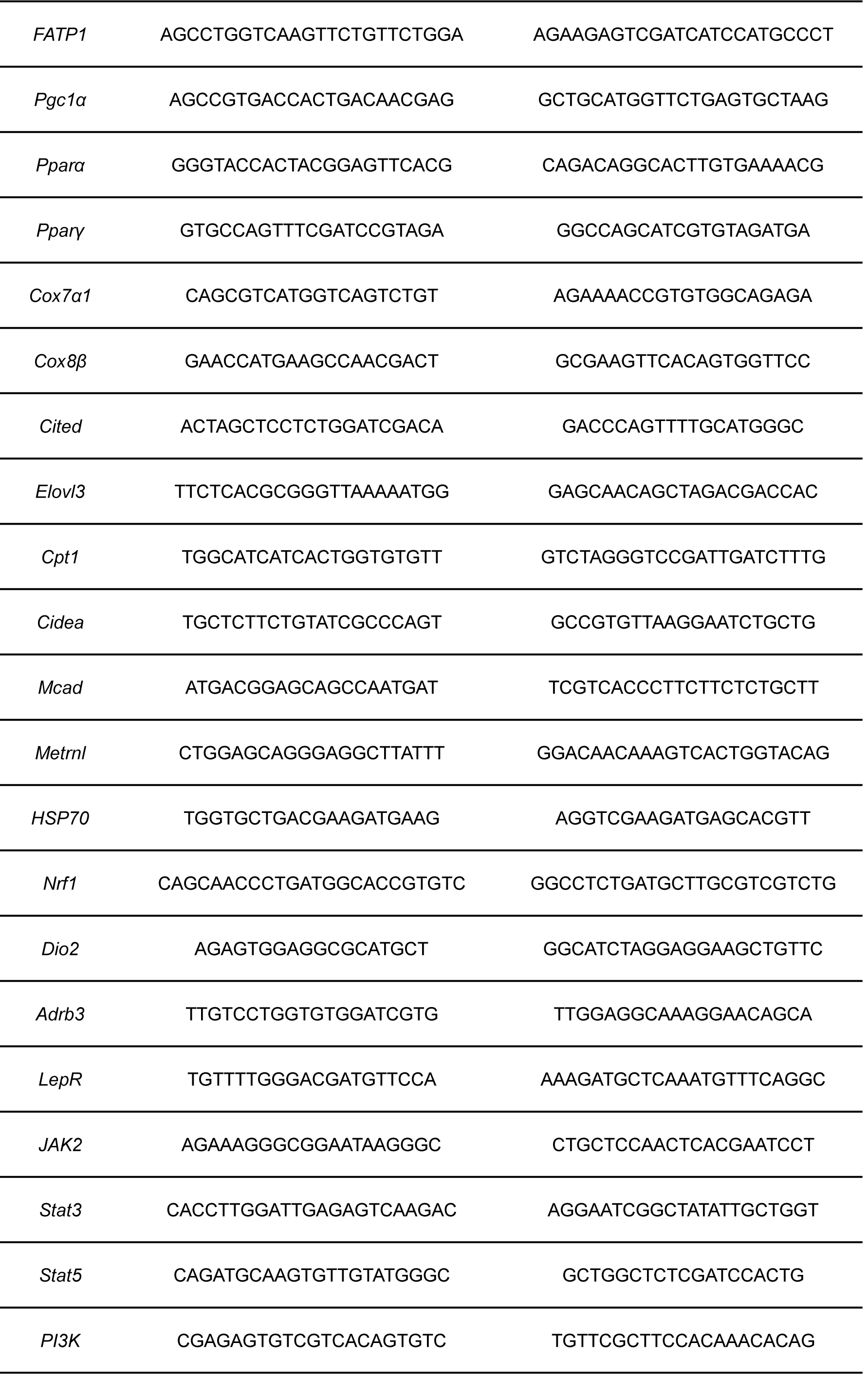
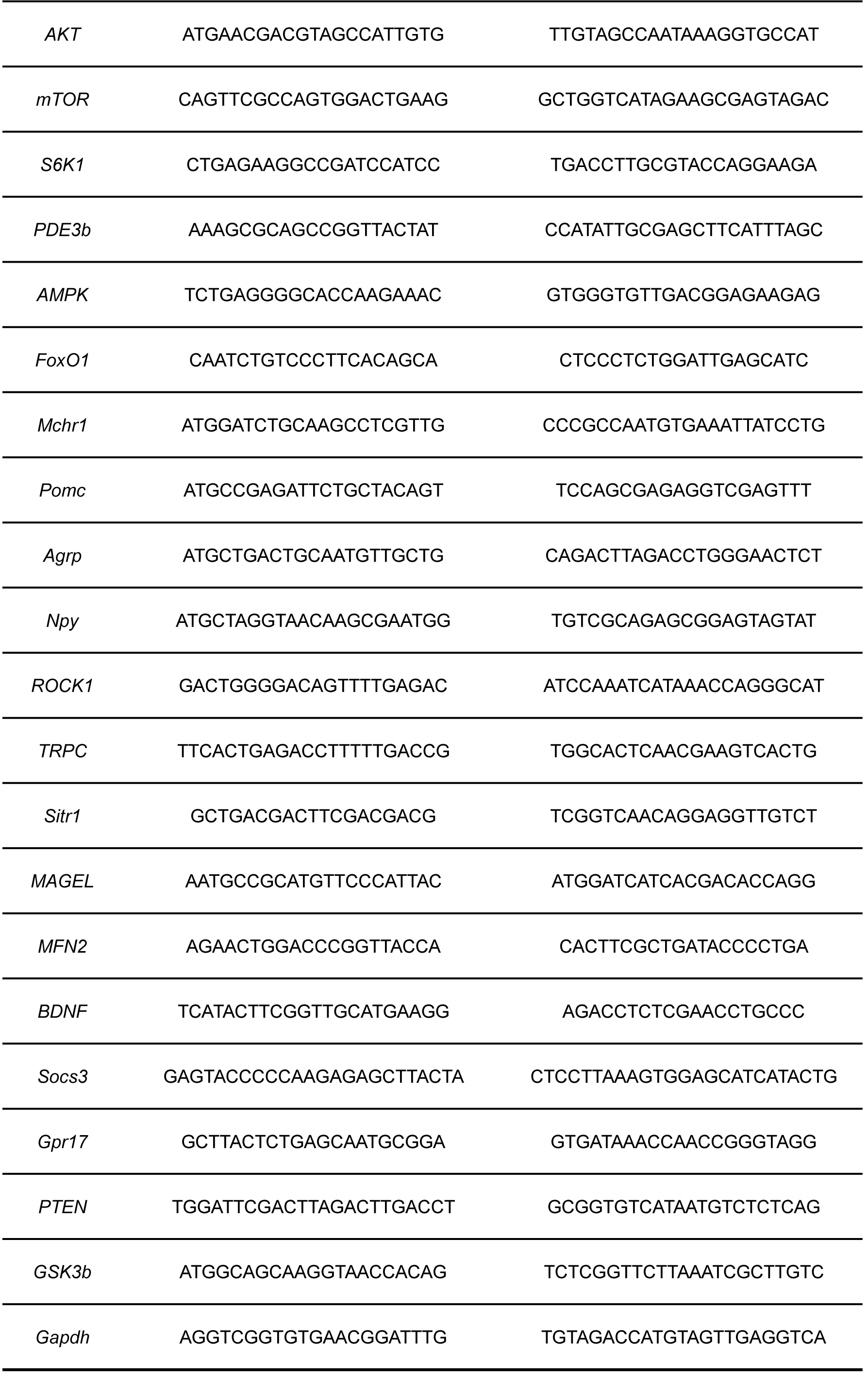
Primers used in the present study

## Notes

### Competing Interest Statement

The authors have declared no competing interest.

